# The vitamin A ester retinyl propionate has a unique metabolic profile and higher retinoid-related bioactivity over retinol and retinyl palmitate in human skin models

**DOI:** 10.1101/2020.09.26.315036

**Authors:** Donald L. Bjerke, Rui Li, Jason M. Price, Roy L.M. Dobson, MyriamRubecca Rodrigues, ChingSiang Tey, Laura Vires, Rachel L. Adams, Joseph D. Sherrill, Peter B. Styczynski, Kirsty Goncalves, Victoria Maltman, Stefan Przyborski, John E. Oblong

## Abstract

Human skin is exposed daily to environmental stressors, which cause acute damage and inflammation. Over time this leads to morphological and visual appearance changes associated with premature aging. Topical vitamin A derivatives such as retinol (ROL), retinyl palmitate (RPalm), and retinyl propionate (RP) have been used to reverse these changes and improve the appearance of skin. This study investigated a stoichiometric comparison of these retinoids using *in vitro* and *ex vivo* skin models. Skin biopsies were treated topically to compare skin penetration and metabolism. Treated keratinocytes were evaluated for transcriptomics profiling and hyaluronic acid (HA) synthesis and treated 3D epidermal skin equivalents were stained for epidermal thickness, Ki67, and filaggrin. A retinoic acid receptor-alpha (RARα) reporter cell line was used to compare retinoid activation levels. Results from *ex vivo* skin found that RP and ROL have higher penetration levels compared to RPalm. RP is metabolized primarily into ROL in the viable epidermis and dermis whereas ROL is esterified into RPalm and metabolized into the inactive retinoid 14-hydroxy-4,14-*retro*-retinol (14-HRR). RP treatment yielded higher RARα activation and HA synthesis levels than ROL whereas RPalm had a null effect. In keratinocytes, RP and ROL stimulated similar gene expression patterns and pathway theme profiles. In conclusion, RP and ROL show a similar response directionality whereas RPalm response was inconsistent. Additionally, RP has a consistently higher magnitude of response compared with ROL or RPalm.

## 1 INTRODUCTION

The skin is the largest organ of the human body and one of its primary functions is to provide protection from damaging external stressors such as solar UV radiation, carbon emissions, and pollution. Cumulative exposure to these damaging stressors leads to structural and functional changes that manifest as older aged skin appearance and has been ascribed as premature aging. Of the stressors, UV radiation exposure is considered the most significant, contributing ∼85% of premature aging and these changes in skin appearance are thus ascribed as photoaging.^[1,2]^

Retinoids represent a class of lipophilic vitamin A derivatives that have been used for decades in cosmetic and pharmaceutical products for photoaging repair as well as for the treatment of acne, psoriasis, ichthyosis, and actinic keratosis.^[3]^ Relative to photoaging, retinoids are known to be critical in maintaining epidermal homeostasis, particularly regulating proliferation and differentiation of keratinocytes and maintaining epidermal thickness.^[4-6]^ Mechanistically, the primary active retinoid form is *trans*-retinoic acid (tRA), which binds to members of the retinoic acid receptor (RAR) family of nuclear hormone receptors and then translocate into the nucleus to activate selective gene expression.^[7,8]^ Retinoids are obtained through the diet in the form of retinol and retinyl esters and as the provitamin in the form of β-carotene which is hydrolyzed into ROL and distributed throughout the body via retinol binding proteins.^[9]^ While the primary route of obtaining vitamin A is via diet and oral absorption, retinoids are also bioavailable when applied topically.^[10]^ Like most eukaryotic cells, epidermal keratinocytes and dermal fibroblasts have the capability to enzymatically convert retinyl esters to ROL and ROL to tRA through the intermediate retinal via an NAD+ dependent oxidative pathway^[11-13]^ where retinoic acid receptors are located in the skin.^[14]^

The impact of topical tRA, ROL, and RPalm on photoaged skin has been previously studied and well documented.^[15-18]^ While ROL has been reported to be efficacious against skin photoaging^[6]^ RPalm has a much lower response, rendering its low irritation profile as moot in terms of being able to have any positive benefits for affecting the appearance of photodamaged skin.^[19]^ This is more than likely due to limited skin penetration and RPalm being a primary storage form for endogenous retinol^[20,21]^, serving as a key regulatory point for maintaining adequate cellular concentrations of retinoids.^[22]^ Moreover, the higher lipophilicity of RPalm (log P 13.8) versus ROL (log P 5.7) and RP (log P 6.7) makes it more likely to remain in the phospholipid bilayers of keratinocytes cellular membrane. RP is a relatively newer retinoid that has been reported to clinically impact photoaged skin with minimal irritation.^[23-25]^ Additionally, besides its overall efficacy and skin tolerability, RP has also been reported to have a better chemical stability profile compared to other esters, thereby increasing its half-life on the skin’s surface during topical delivery.^[26]^

To better understand the bioavailability and the biological activity profile of these topically applied retinoids, we evaluated RP, ROL, and, in some cases, RPalm in human skin models. The evaluations included skin penetration and metabolic fate, gene expression response profiles, retinoid receptor activation, and cellular responses in keratinocytes and 3D epidermal skin equivalents.

## 2 METHODS

### 2.1 Skin biopsy dermal penetration and metabolism

Topical formulations were prepared to take into account the weight differences between RP, RPalm, and ROL so that all were at a stoichiometrically equivalent concentration of 0.3% ROL. The retinoids tested included 0.3% [^14^C]-retinol, 0.359% [^14^C]-retinyl propionate, or 0.55% [^14^C]-retinyl palmitate (Pharmaron, Inc.), and were added to the oil phase pre-mix during preparation of oil-in-water emulsions at the stated concentrations Four samples of full-thickness Caucasian human skin (abdominal) were obtained from three female (ages 34, 40, and 41) and one male donor (age 52) following GLP and OECD 428 guidelines. Skin samples were obtained from NHS Lothian (Edinburgh, Scotland) from patients undergoing cosmetic abdominoplasty. The subcutaneous fat and connective tissue were removed and a split-thickness layer (600-800 μm) was prepared with a dermatome. Human skin samples were used immediately following preparation. The metabolic competence of all fresh skin donors was assessed by measuring the rate of MTT ([3,4, 5-dimethylthiazol-2-yl]-2,5-diphenyltetrazolium bromide) reduction to its formazan metabolite by mitochondrial reductase enzymes. In addition, phenyl acetate hydrolysis was assessed at 0, 4, and 24 hours, to confirm enzyme activity of the skin. Discs of dermatomed skin were obtained and mounted in 6-well static diffusion cells (exposed skin area, 2.27 cm^2^). The receptor fluid was HEPES (25 mM) buffered Hank’s balanced salt solution containing bovine serum albumin (4%, w/v), pH 7.32-7.46. Plates were then placed in a humidified incubator (*ca* 5% CO_2_ and *ca* 32°C) on a rocker plate. Formulations were manufactured with ^14^C-labelled retinol, retinyl propionate and retinyl palmitate at 0.3% retinol equivalent concentrations in an oil in water face cream emulsion. The ^14^C label was positioned on the last carbon of the alkyl side chain before the alcohol or ester moiety. The retinoid formulations (5 mg/cm^2^) were applied to each diffusion and receptor fluid was sampled at 8 and 24 hours after dosing. At the end of the 24 hours, the skin was removed from the diffusion cell and the amount of retinoid remaining in the stratum corneum, viable epidermis, dermis and receptor fluid was determined. Skin samples were tape stripped fifteen times to remove the stratum corneum. Each tape strip was placed into a scintillation vial to measure total ^14^C-labeled retinoid. The epidermis was separated from the dermis using a scalpel. The epidermis and dermis samples were frozen in liquid nitrogen, stored at −80°C, and subsequently prepared for extraction by biopulverisation after freezing. The skin homogenates were then extracted in ethanol. Aliquots of the ethanol extracts were removed for analysis by liquid scintillation counting and the remaining sample was analyzed by high performance liquid chromatography, with on-line mass spectrometry and radiochemical detection (HPLC-MS-RAD). All sample extracts were stored at −80°C until analysis. Samples analyzed for retinoid metabolites included the viable epidermis and dermis. Due to small sample volumes, low concentrations of retinoids in the receptor fluid, and the viscous nature of the samples, the receptor fluid was not suitable for metabolite analysis. Epidermal and dermal extracts were dried and reconstituted in methanol:propan-2-ol (7:3 v/v). Procedural recoveries for preparation of the epidermis and dermis extracts were in the ranges of 50-84% and 42-92%, respectively, which were deemed acceptable due to low radioactivity and low sample volumes. For sample analysis, a Thermo Q-Exactive+ MS instrument, operated in heated electrospray ionization (HESI) and positive/negative switching modes, was employed in conjunction with a Shimadzu Nexera Modular HPLC system and LabLogic β-RAM Model 5 RAD. Approximately 25% of HPLC effluent was diverted to the MS using an inline splitter, with the remainder sent to the RAD. HPLC conditions included a Phenomenex Security Guard C18 precolumn and Phenomenex Prodigy ODS2 column, using 2 mM ammonium acetate in water:methanol (1:9 v/v) and 2 mM ammonium acetate in water:propan-2-ol (1:9 v/v) gradient mobile phase at a flow rate of 1.0 mL/min. These conditions afforded retention and resolution of all three retinoids (RP, ROL, RPalm) and metabolite identification using a single HPLC method. HESI-MS detection of ROL RP and RPalm all resulted in formation of a characteristic fragment ion, observed in the full-scan mass spectrum. Detection of this positive ion fragment, *m/z* 269.2264, formed by neutral-loss of H_2_O, propionate and palmitate moieties from protonated ([M+H]^+^) ROL, RP and RPalm, respectively, and correlated with HPLC retention times for ROL, RP and RPalm reference standards served to verify parent compound identities within each extract. The identities of incurred ^14^C-containing metabolites were investigated using HPLC-MS-RAD by comparison of retention times, accurate mass measurements, and expected isotope and fragmentation patterns, with those obtained from control samples and authentic synthetic reference standards. Reference standards for potential retinol metabolites including tRA, 14-hydroxy-4,14-*retro*-retinol (14-HRR), dehydroretinol, anhydroretinol, RPalm, retinyl oleate and retinyl stearate were additionally analyzed to specifically investigate whether any of these could be assigned to the observed ^14^C-labeled metabolites. In addition to facilitating metabolite identification, the HPLC-RAD profiles enabled relative quantitation of ^14^C-labeled species within each extract. Control samples were run for all donors. Due to retinoids being light sensitive, procedures were carried out, where possible, under yellow light in order to avoid photo-degradation.

### 2.2 Microarray analysis

Human telomerised (hTERT) keratinocytes (obtained as a kind gift from Dr. Jerry Shay, University Texas Southwestern) were maintained in KBM-Gold Keratinocyte Growth Medium BulletKitTM (Lonza) according to manufacturer’s protocol. Cells at passage 3 were left untreated or treated with 100 nM RP or 100 nM ROL (Sigma) for 24 hours. Samples were collected, processed, and subjected to microarray profiling on the GeneTitan U219 array platform (Affymetrix, Santa Clara, CA) as previously described.^[27]^ Data were analysed using GeneSpring version 14.8 (Agilent Technologies, Santa Clara, CA). Quartile normalization and Plier summarization was performed on all probe sets. Probe sets with low expression (< 20^th^ percentile across all samples within any one age group) were removed and a one-way ANOVA with Tukey post-test was performed to identify statistically significant probe sets (Benjamini-Hochberg corrected *p-*value < 0.05) across the 3 treatment groups (no treatment, 100 nM RP, 100 nM ROL). Hierarchical clustering was performed using Euclidean distance.

### 2.3 Quantitative PCR Analysis

hTERT keratinocytes were cultured with Epilife media supplemented with human Keratinocyte Growth Supplement and Gentamicin/Amphotericin B 500X (Thermo Fisher Scientific, USA). Cells grown to passage 3 were treated for 24 hr with vehicle (DMSO) or 100 nM, 250 nM,1 uM, or 5 uM ROL or RP (Sigma-Aldrich, USA). Cell lysates were collected and RNA was isolated using the Biomek FxP and the RNAdvance Tissue Isolation kit (Beckman Coulter, USA, p/n A32646). The resulting RNA was quantified using the Nandrop 8000 (Nanodrop, ND-8000). cDNA was generated using 500 ng of TotalRNA and Applied Biosystems High Capacity cDNA with Reverse Transcription kit (Applied Biosystems p/n 4368814). cDNA, assays, and dilutions of PrimeTime GeneExpression MasterMix (IDT, p/n 1055771) were plated onto a Wafergen MyDesign SmartChip (TakaraBio, p/n 640036) using the Wafergen Nanodispenser. The chip was then loaded into the SmartChip cycler and qPCR performed using the following conditions: Hold Stage - 50°C for 2 minutes, 95°C for 10 minutes; PCR Stage - 95°C for 15 seconds (40 cycles), 60°C for 1 minute. Relative expression values of target genes were normalized to the geometric mean of 4 housekeeping genes (*ACTB, B2M, GAPDH*, and *PPIA*) and fold changes over vehicle-treated cells was evaluated for significance using a Student’s t-test. A complete list of the target genes analysed and qPCR data can be found in Supplementary Table 2.

### 2.4 RARα reporter activation measurements

HEK293 RARα reporter cells (BPS Bioscience, San Diego, CA) were grown in assay medium (DMEM, high glucose, no phenol red (LifeTech, Singapore), 10% coal-stripped FBS (GIBCO,), 1x penicillin streptomycin (LifeTech, Singapore), 4mM L-glutamine (Sigma-Aldrich, Singapore)) for 24 hours. Harvested cells were normalized to ∼330,000 cells per ml and 90 □l was seeded into each well for a final cell count of ∼30,000 per well. Retinoid stock solution were at 100 mM in DMSO and serially diluted for final concentration. Cells were treated with 15.6, 62.5, 250, 1000, or 4000 nM RP, ROL, or RPalm for 24 hours. Cell viability was measured by CellTiter as per manufacturer’s instructions (Promega, USA). Luciferase activity was measured by Bio-Glo as per manufacturer’s instructions (Promega, USA). CellTiter and Bio-Glo luminescence measurements were performed using a Cytation 3 Imaging Reader.

### 2.5 Hyaluronic acid synthesis and staining

hTERT keratinocytes were maintained in KGM^™^ Gold^™^ Basal Medium (Lonza, Singapore) supplemented with KGM^™^ Gold^™^ SingleQuots^™^ supplements (Lonza, Singapore) and incubated at 37°C, 5% CO_2_ in a humidified environment. Cells grown to passage 3 were treated for 48 hr with vehicle (DMSO), or 6.25 nM, 12.5 nM, 25 nM, 50 nM, or 100 nM ROL or RP, or 100 nM RPalm. Cell culture supernatant was collected and stored at −20°C for quantification of HA. Cells were fixed with 4% formaldehyde (Sigma-Aldrich, Singapore) in DPBS (Gibco, Singapore) for 15 minutes at room temperature, before staining with NucBlue^™^ (Invitrogen, Singapore) according to the manufacturer’s instructions for automated cell count by BioTek Cytation 3 Imaging Reader. The quantification of HA was done with the hyaluronan quantikine ELISA kit (R&D Systems, Singapore) according to the manufacturer’s instructions. Optical density of each well was determined by using BioTek Cytation 3 Imaging Reader at 450 nm, with wavelength correction at 540 nm.

For HA staining, cells were seeded on coverslip placed in wells of a 6-well plate for 24 hours, before treatment for 48 hours with vehicle (DMSO) or 100 nM RP. Cells were fixed for 10 minutes with cold (−20°C) methanol (Sigma-Aldrich, Singapore), blocked for 2 hours with 1% BSA (Sigma-Aldrich, Singapore) in DPBS (Gibco, Singapore) at 4°C. Biotinylated hyaluronic acid binding protein (Sigma-Aldrich, Singapore) were diluted in blocking buffer and incubated with samples overnight at 4°C. Samples were then washed three times with cold DPBS (Gibco, Singapore) and incubated with Alexa Fluor^™^ 594-conjugated steptavidin (Invitrogen, Singapore) diluted in blocking buffer for 2 hours at room temperature in the dark. Samples were then washed three times with cold DPBS (Gibco, Singapore) and mounted using ProLong™ Glass Antifade Mountant with NucBlue™ Stain (Hoechst 33342, Invitrogen, Singapore). Fluorescent images were generated using the Zeiss LSM 710 confocal microscope with ZEN software.

### 2.7 Construction and treatment of 3D epidermal skin equivalents with RP and ROL

*In vitro* 3D epidermal skin equivalents were generated as previously described.^[28]^ Human neonatal keratinocytes (HEKn, ThermoFisher Scientific, Loughborough, UK) were maintained as per manufacturer’s instructions. Cells were maintained in Epilife^®^ medium (ThermoFisher Scientific, Loughborough, UK) supplemented with Human Keratinocyte Growth Supplement (HKGS) and incubated at 37°C, 5% CO_2_ in a humidified environment. Briefly, HEKn were seeded onto a collagen I (ThermoFisher Scientific, Loughborough, UK) coated Millicell^®^ culture membrane (Merck Millipore, Watford, UK) in submerged culture, low calcium conditions for 2 days. Cultures were then moved to the air-liquid-interface (ALI) in high calcium conditions for 10 days. At day 10, ALI models were treated with retinoids. RP (Sigma-Aldrich, St Louis, USA) and ROL (Sigma-Aldrich, St Louis, USA) were added to culture medium in a vehicle of dimethyl sulfoxide (DMSO, Sigma-Aldrich, St Louis, USA) at a final concentration of 0.01 μM or 10 μM. Culture medium was replaced with fresh retinoid containing medium and models were harvested after 3 days of treatment.

### 2.8 Histology and immunofluorescence staining of 3D epidermal skin equivalents treated with RP or ROL

3D epidermal skin equivalents were harvested by fixation in 4% paraformaldehyde (Sigma-Aldrich, St Louis, USA) for 2 hours at room temperature. Samples were then dehydrated through a series of ethanol washes (30% - 100%), incubated in Histo-Clear II (Scientific Laboratory Supplies, Nottingham, UK) followed by a 1:1 mix of Histo-Clear II and paraffin wax, before being embedded in plastic moulds (CellPath, Newton, UK). 3D epidermal skin equivalents were sectioned transversely using a microtome (Leica RM2125RT) at 5 μm. Sections were transferred to charged microscope slides (ThermoFisher Scientific, Loughborough, UK) for analysis.

Samples of 3D epidermal skin equivalents were deparaffinised in Histo-Clear I (Scientific Laboratory Supplies, UK) and rehydrated through a series of ethanol baths. Samples were then incubated in Mayer’s haematoxylin (Sigma-Aldrich, St. Louis, USA) for 5 minutes and nuclei were blued in alkaline ethanol for 30 seconds. Slides were then dehydrated through a series of ethanol washes before being incubated with eosin (Sigma-Aldrich, St. Louis, USA) for 30 seconds and further dehydrated. Samples were then cleared in Histo-Clear I prior to mounting with Omnimount (Scientific Laboratory Supplies, UK). Histology images were generated using Leica ICC50 high definition camera and a Brightfield Leica microscope. For immunofluorescence staining, antigen retrieval was conducted on deparaffinised 3D epidermal skin equivalents by incubation at 95°C in citrate buffer (Sigma-Aldrich, St. Louis, USA) for 20 mins. Samples were permeabilised and blocked for an hour in a solution of 20% neonatal calf serum (Sigma-Aldrich, St. Louis, USA) in 0.4 % Triton X-100 (Sigma-Aldrich, St. Louis, USA) in phosphate buffered saline (PBS). Primary antibodies targeting Ki67 ab16667 (Abcam, Cambridge, UK) and Filaggrin ab17808 (Abcam, Cambridge, UK) were diluted in blocking buffer and incubated with samples overnight at 4°C. Samples were then washed three times in PBS and incubated with secondary antibodies diluted in blocking buffer for an hour at room temperature: anti-rabbit Alexa Fluor^®^ 488 nm (ThermoFisher Scientific, Loughborough, UK) and anti-mouse Alexa Fluor^®^ 594 nm (ThermoFisher Scientific, Loughborough, UK). Samples were then washed in PBS three times and mounted using Vectasheild with DAPI Hardset Mounting Medium (Vector Laboratories, Peterborough, UK). Fluorescent images were generated using the Zeiss 880 confocal microscope with ZEN software. For image analysis, measurements taken from images were obtained using Image J software (https://imagej.nih.gov). Briefly the scale was set and the line tool selected, to take measurements of the viable epidermis for quantification. Image J was also used to count the number of Ki67 positive nuclei using the multipoint tool and percentage positive nuclei was then calculated.

## 3 RESULTS

### 3.1 Topically applied RP and ROL penetrate to the viable epidermis and dermis at similar levels, but after 24 hours RP yields higher ROL in these skin compartments as compared to topically applied ROL and RPalm

To evaluate the skin penetration and metabolic fate of ROL, RP, and RPalm, freshly collected human split-thickness skin samples were treated with 0.3% retinol weight equivalent concentrations of radiolabelled retinoids in an oil-in-water emulsion (0.3% [^14^C]-retinol, 0.359% [^14^C]-retinyl propionate, or 0.55% [^14^C]-retinyl palmitate). After 24 hours of treatment, total recovery of radioactivity was within 90-110%, and the majority of the applied retinoids applied were washed off (87-89%). The amount of retinoid in the tape-stripped stratum corneum ranged from 6 to 7% (data not shown). The amount absorbed into the receptor fluid (mean ± SD) over 24 hours was 0.92 ± 0.31% for ROL, 0.31 ± 0.15% for RP, and 0.09 ± 0.08% for RPalm (Table 1). The amount in the viable epidermis after 24 hours was 0.85 ± 0.52% for ROL, 0.64 ± 0.64% for RP, and 0.28 ± 0.25% for RPalm. The amount in the dermis after 24 hours was 0.33 ± 0.23% for ROL, 0.15 ± 0.09% for RP, and 0.04 ± 0.02% for RPalm. In units of µg equivalents/cm2, delivery to the viable epidermis and dermis was 0.19 ± 0.10, 0.14 ± 0.11, and 0.09 ± 0.07 for ROL, RP, and RPalm, respectively. Dermal delivery, which includes amount in the viable epidermis, dermis and receptor fluid was 2.10 ± 0.78% for ROL, 1.09 ± 0.59% for RP, and 0.41 ± 0.25% for RPalm. In units of µg equivalents/cm^2^, dermal delivery was 0.32 ± 0.12, 0.20 ± 0.11, and 0.11 ± 0.07 for ROL, RP, and RPalm, respectively. The values represent 4 replicates per donor with a total of 4 donors. In addition to physical delivery of the retinoids, we also characterized the metabolic fate of the ^14^C-labeled forms in the viable skin layers. The metabolic competence of all fresh skin donors was confirmed via MTT reduction and esterase activity via phenyl acetate hydrolysis (data not shown). After topical treatment for 24 hours, the individual skin compartments were separated and homogenized, extracted in ethanol and analyzed by HPLC-MS-RAD (representative HPLC-RAD traces can be seen for each retinoid in Supplemental Figure 1). ROL treatment showed the major metabolite was the inactive analogue 14-hydroxy-4,14-*retro*-retinol (14-HRR) in epidermis (39.3 ± 12.3% of the regions of interest) and dermis (32.7 ± 9.1% of the regions of interest) followed by the parent compound ROL (12.1 ± 2.3% and 6.4 ± 2.0%, respectively) and the storage form RPalm (4.7 ± 1.6% and 3.1% (single value, found in 1 of 4 donors), respectively) (Table 2). In contrast, RP application delivered more parent compound to the epidermis (53.4 ± 17.1%) and dermis (42.6 ± 11.4%) and the ester hydrolysis product ROL (24.1 ± 12.7% and 12.7 ± 6.0%, respectively). This is double the proportion of ROL present after 24 hours when compared to when equimolar ROL is applied to the skin (Table 2). This translates into 0.12% of the ROL dose delivered to viable skin as ROL, versus 0.17% of the RP dose delivered as ROL. Further, at 24 hours, 0.40% of the RP dose remains in viable skin as not-yet-metabolized parent, potentially as a source of additional ROL through subsequent hydrolytic release. This suggests a 0.57% total ROL delivery potential, when dosed as RP (almost 5x greater than achieved via ROL treatment). Also, inactive 14-HRR and the storage form RPalm were not detected in viable epidermis and dermis after 24 hours of topical exposure to RP. Following RPalm topical treatment, only the parent compound was found in the epidermis (83.9 ± 2.7%) and dermis (55.1 ± 19.2%), with no ROL or 14-HRR detected.

**Table 1.**
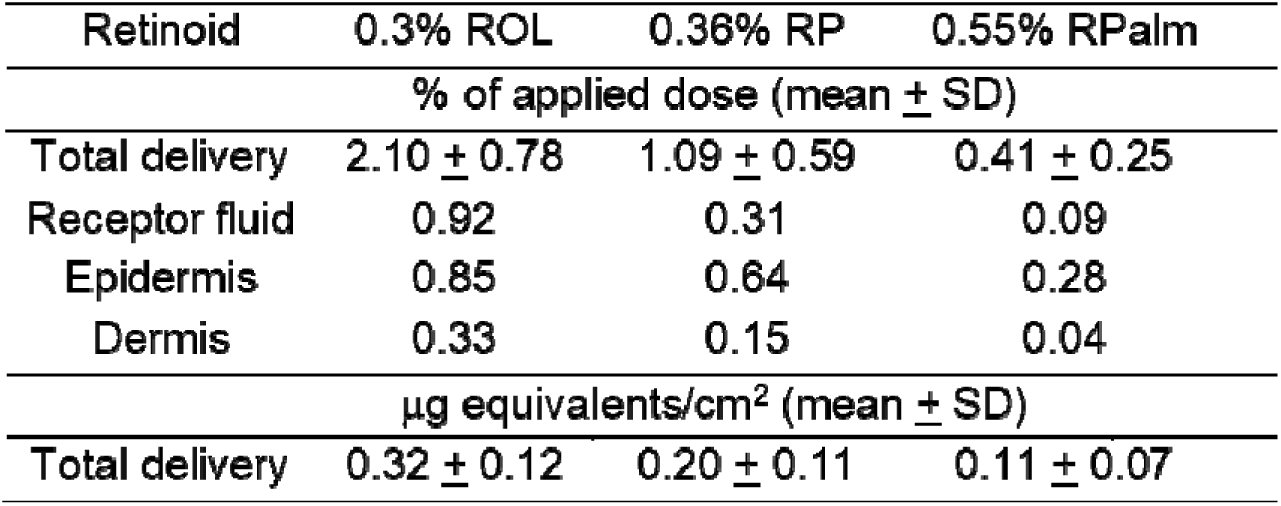
Skin delivery of retinoids following 24 hours of topical treatment with ROL, RP, or RPalm

| Retinoid | 0.3% ROL | 0.36% RP | 0.55% RPalm |
| --- | --- | --- | --- |
| % of applied dose (mean ± SD) | | | |
| Total delivery | 2.10 ± 0.78 | 1.09 ± 0.59 | 0.41 ± 0.25 |
| Receptor fluid | 0.92 | 0.31 | 0.09 |
| Epidermis | 0.85 | 0.64 | 0.28 |
| Dermis | 0.33 | 0.15 | 0.04 |
| μg equivalents/cm<sup>2</sup> (mean ± SD) | | | |
| Total delivery | 0.32 ± 0.12 | 0.20 ± 0.11 | 0.11 ± 0.07 |

**Table 2.** Retinoid metabolites from skin biopsies following 24 hours of topical treatment with RP, ROL, or RPalm

| Retinoid | Epidermis (% ROI ± SD) | | | Dermis (% ROI ± SD) | | |
| --- | --- | --- | --- | --- | --- | --- |
|  | 0.3% ROL | 0.36% RP | 0.55% RPalm | 0.30% ROL | 0.36% RP | 0.55% RPalm |
| ROL | 12.1 ± 2.3 | 24.1 ± 12.7 | ND | 6.4 ± 2.0 | 12.7 ± 6.0 | ND |
| 14-HRR | 39.3 ± 12.3 | ND | ND | 32.7 ± 9.1 | ND | ND |
| RPalm | 4.7 ± 1.6 | ND | 83.9 ± 2.7 | 3.1* | ND | 55.1 ± 19.2 |
| RP | ND | 53.4 ± 17.1 | ND | ND | 42.6 ± 11.4 | ND |

ROI = region of interest in radiochromatogram; ND – not detected; \* detected in only one donor sample, value reflects single donor only.

**Figure 1.**
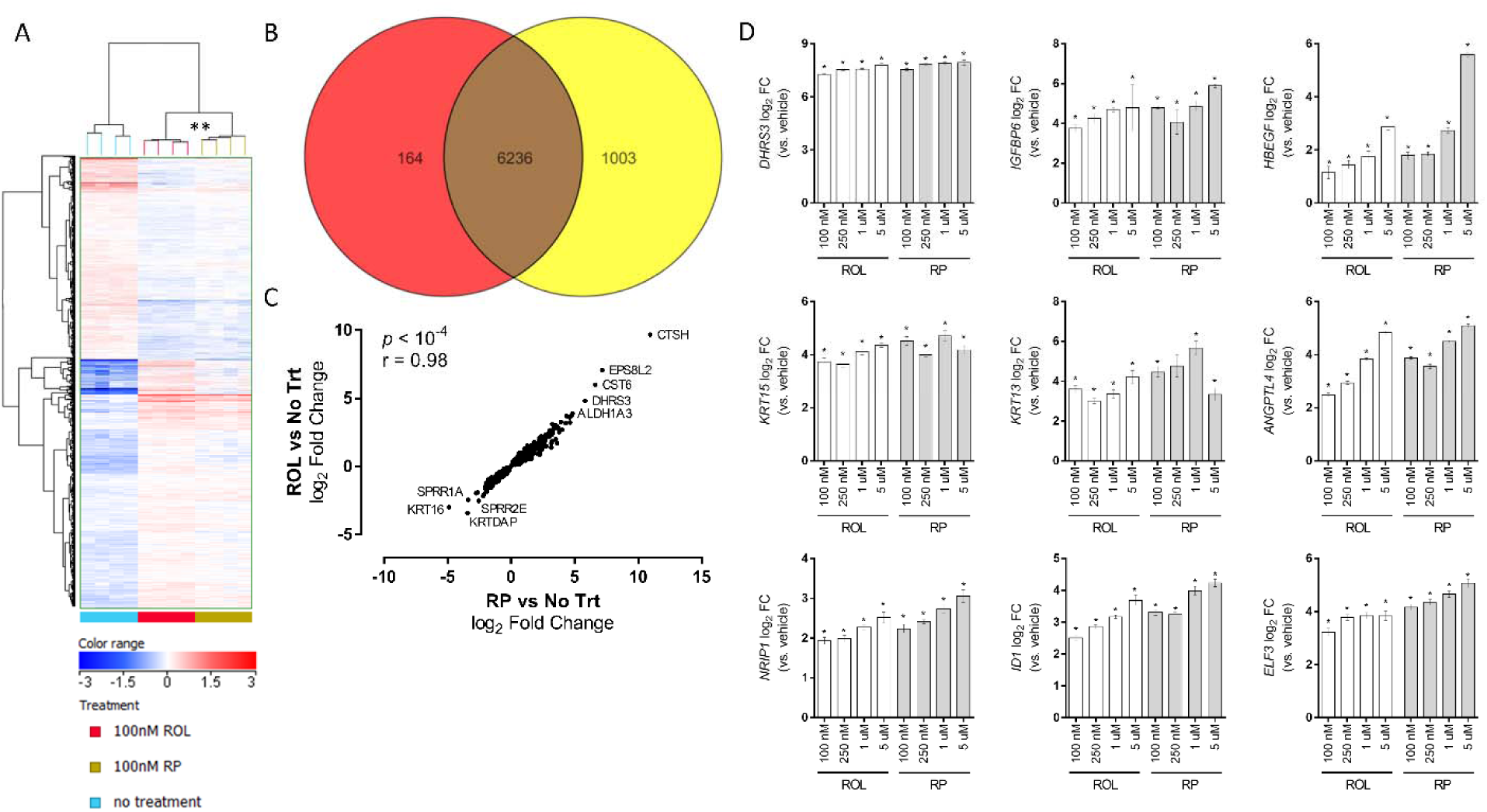
Gene expression profiles comparing ROL and RP show equivalent responses on expression patterns and pathway impact. A, Hierarchical clustering of significant probe sets (n = 7,418; ANOVA corrected *p*-value < 0.05) and replicate samples across treatment groups (RP, ROL, or no treatment (No Trt)). Color range of heatmap shows log2 expression values. B, Venn diagram showing the number of common significant probe sets (n = 6,236) between the ROL (red) and RP (yellow) signatures vs. no treatment (No Trt). C, Pearson correlation of log2 fold changes of common significant probe sets (n = 6,236) between RP vs. No Trt (x-axis) and ROL vs. No Trt (y-axis). D, Independent replication of ROL and RP target genes by qPCR. hTERT-KCs were treated for 24 hours with 100 nM – 5 uM ROL or RP. Shown are the log2 fold changes (FC) of select retinoid target genes (vs. untreated) across multiple doses. Student’s t-test, \**p* < 0.05. n = 12 per treatment group. Error bars represent log2 SEM.

### 3.2 ROL and RP have overall equivalent stimulatory effects on gene expression patterns in treated hTERT keratinocytes

To determine what similarities there are between ROL and RP on stimulating gene expression patterns *in vitro* in keratinocytes, we performed microarray analyses on hTERT keratinocytes treated with 100 nM ROL or RP for 24 hours. We identified a total of 7,418 probe sets that were differentially expressed across treatment groups (one-way ANOVA corrected, \*\**p* < 0.05) with a strikingly similar expression pattern between ROL and RP compared to control (Figure 1A). The complete list is in Supplementary Table 1. Indeed, 6,236 significant probe sets were common to both the ROL and RP signatures (Figure 1B). Moreover, these shared probe sets showed a near-linear correlation in fold change vs. no treatment (Pearson r = 0.98, *p* < 10^−4^), with the RP signature showing a greater dynamic range compared to the ROL signature (Figure 1C).

We subsequently validated the transcriptional changes observed by microarray and the enhanced regulation by RP compared to ROL of 48 target genes by qPCR in an independent experiment. Here, we observed complete conservation of the regulatory pattern for all 48 target genes (40 upregulated, 8 downregulated) by both RP and ROL in accordance with the microarray results. In addition, the majority of the target genes were regulated in a dose-responsive manner, with RP eliciting a numerical greater effect than ROL compared to control on 31 target genes (Supplementary Table 2). Bar charts demonstrating the response profiles of 9 select retinoid target genes across the different doses and treatments are shown in Figure 1D.

### 3.3 RP stimulates a higher RARα activation and increase in HA synthesis than ROL and RPalm in a dose dependent manner

A RARα reporter (Luc)-HEK293 cell line that contains a luciferase gene under the control of retinoic acid response elements stably integrated into HEK293 cells along with full length human RARα was used to quantitate the effect of ROL, RPalm, and RP on inducing expression of the luciferase gene. This reporter system requires the presence of tRA and thus is an indirect measure of enzymatic conversion of ROL, RPalm, and RP into tRA once inside the cells (Figure 2A). A dose response of tRA was used to establish the luciferase signal range (data now shown). At 15.6 nM tRA induced luciferase activity that was 463% higher than vehicle control (Figure 2B). ROL, RPalm, and RP were tested at 15.6, 62.5, 250, 1000, and 4000 nM and signal response was normalized to vehicle control. While both ROL and RP showed a dose dependent increase in luciferase activity, RP had a 20-180% greater signal intensity than ROL across the doses tested, suggesting a higher concentration of synthesized tRA. Interestingly, RPalm did not stimulate a response and in fact showed significant inhibition at 250, 1000, and 4000 nM that ranged up to 44% compared to vehicle control.

**Figure 2.**
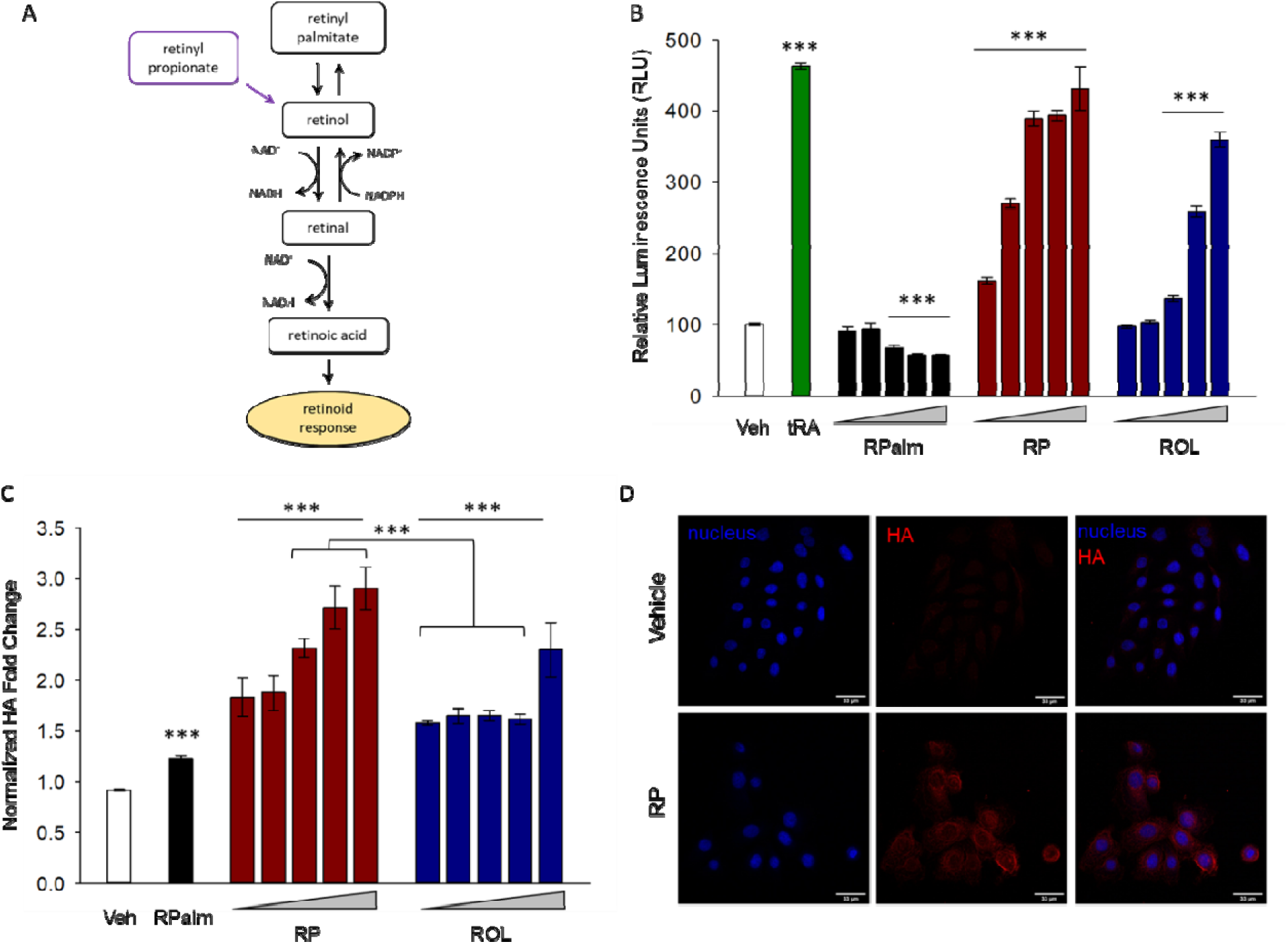
RARα activation and HA synthesis levels are increased by RP to a higher level than by ROL and RPalm. A, Schematic representation of the enzymatic conversion of RP, RPalm, and ROL to the active form of tRA. B, HEK293 cells containing a stable RARα reporter construct were treated for 24 hours with 15.6 tRA and 15.6, 62.5, 250, 1000, and 4000 nM of RPalm, RP, or ROL (dose range represented by increasing triangle). Luciferase activity was quantitated, normalized to control cells and signal compared to vehicle control treatment (Student’s t-test, \*\*\**p*<0.001). C, HA production in hTERT keratinocytes was measured after treatment for 48 hours with ROL or RP at 6.25, 12.5, 25, 50, and 100 nM and with RPalm at 100 nM. Data was normalized to control cells (without treatment, Student’s t-test, \*\*\**p*<0.001). D, HA and nuclei staining using biotinylated-HA binding protein (red) and Hoechst 33342 (blue), respectively, in keratinocytes treated with 100 nM RP for 48 hours.

To further understand the RAR mediated response, we treated keratinocytes with the retinoids and quantitated hyaluronic acid (HA) synthesis levels. HA was selected since it’s known that tRA bound RAR complexes can activate an RAR response element upstream of hyaluronic acid synthase 2 transcription start site in keratinocytes.^[29]^ Additionally, RP and ROL both increased expression of hyaluronic acid synthase 3 (HAS3) in hTERT keratinocytes (Supplementary Table 2). As a positive control, tRA at 100 nM stimulated a 3.5-fold increase in HA secreted levels compared to vehicle control (data not shown). All retinoids at all concentrations tested showed a significant increase in secreted HA levels compared to vehicle control (Figure 2C, \*\*\**p*<0.001). Additionally, RP and ROL at 6.25, 12.5, 25, 50, or 100 nM showed a significant increase in HA over 100 nM RPalm (\*\*\**p*<0.001). RP elicited a more consistent dose response across 6.25, 12.5, 25, 50, and 100 nM. In contrast, there was no dose response effect by ROL across 6.25, 12.5, 25, and 50 nM doses but did show an increase at 100 nM. RP significantly increased HA synthesis when compared to ROL by 40% at 25 nM and by 68% at 50 nM (\*\*\**p*<0.001). At 100 nM, RP was numerically higher by 26% than ROL but was not statistically significant (*p*=0.07). We also visualized an amplification by RP of pericellular bound HA on keratinocytes by staining with a biotinylated HA binding protein and a fluorochrome-tagged streptavidin fluorescent probe (Figure 2D, red color) as well as nuclear staining with Hoechst 33342 (Figure 2D, blue color).

### 3.4 RP has a higher response than ROL on epidermal thickening and Ki67 immunostaining in 3D epidermal skin equivalents

Previously characterised 3D epidermal skin equivalents were used to assess the biological impact of ROL and RP.^[24]^ Retinoid compounds were added to the culture media of models for three days which were fed from beneath and cultured at the air-liquid interface. Histological analysis (Figure 3A) reveals the epidermis was well-structured and viable with all treatment regimes. The epidermis, however, appeared thicker when treated with ROL and RP compared with the vehicle control. Quantification of epidermal thickness confirmed this observation, with ROL treatment resulting in a 21% increase in epidermal thickness compared to vehicle, and RP treatment resulting in a 44% and 16% thicker epidermis than the vehicle and ROL treated samples, respectively (Figure 3B). Total nucleated cells were counted per field of view and both RP and ROL respectively increased cell count by 22% and 107% over vehicle control (Figure 3C). RP increased cell count by 68% over ROL. Similarly, treatment with both retinoids also induced a significant increase in the number of proliferative cells in the epidermis as detected by Ki67 immunoreactivity (green, Figure 3D). The percentage of Ki67 positive cells was increased by 150% by ROL treatment. RP treatment increased the proliferative cell population by 350% and 80% compared with the vehicle control and the ROL samples, respectively (Figure 3E). We also stained for filaggrin, a terminal differentiation marker, which was detected in the vehicle treated epidermis at low levels (red, Figure 3D). An increase in filaggrin positive staining was observed with both ROL and RP treatment, however this was not quantified (data not shown). This shows that both RP and ROL treatments are having a positive effect in this model and that RP has a greater biological effect than ROL on the epidermis *in vitro* in terms of both epidermal thickness and proliferative capacity.

**Figure 3.**
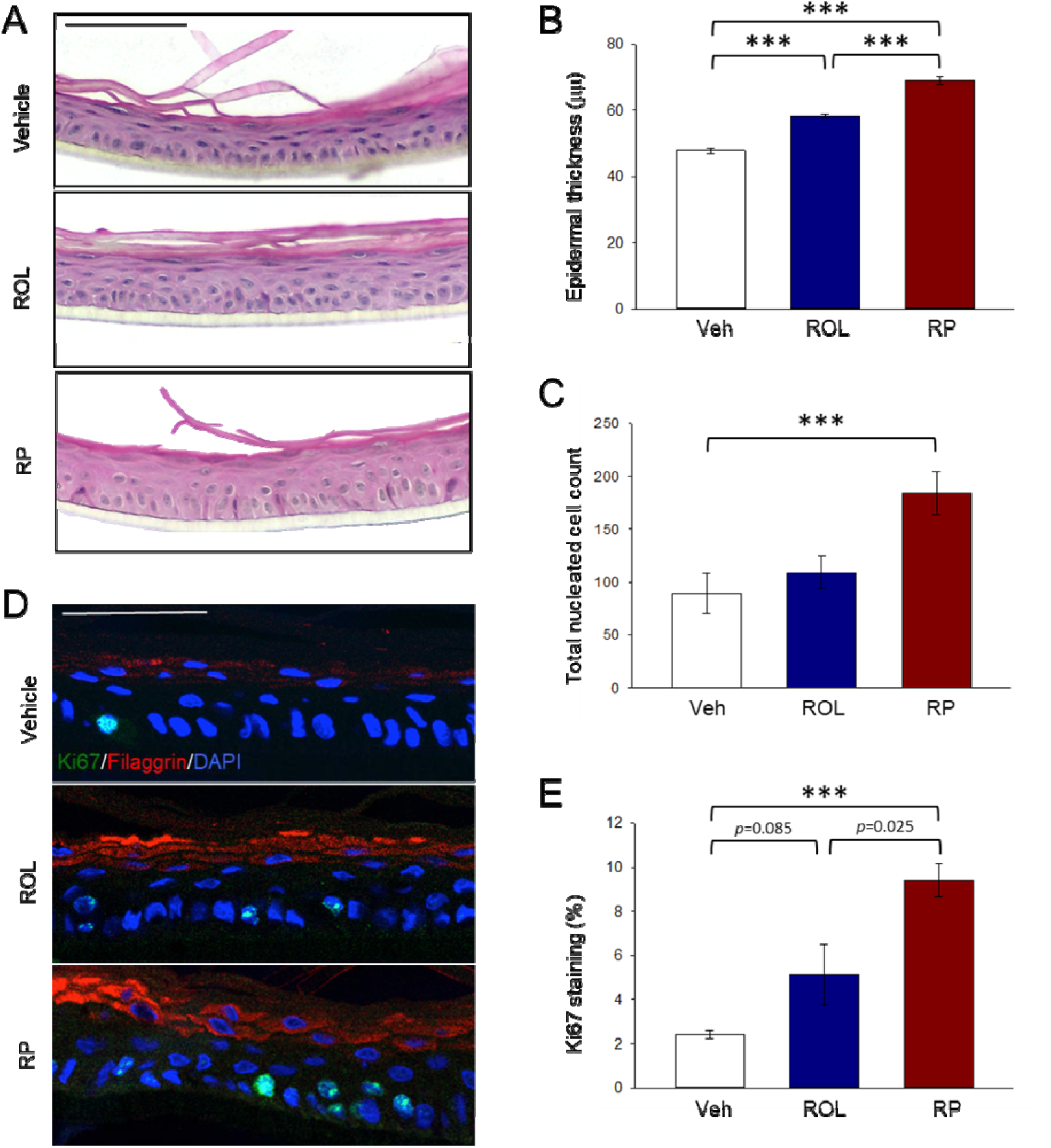
RP shows a higher stimulation than ROL for increasing epidermal thickness and Ki67 staining in 3D epidermal equivalents following retinoid treatment. 3D epidermal equivalent models were treated with 10 μM of ROL or RP for 3 days and then processed for histological and immunological analysis. A, B and C, Epidermal thickening was visualized and total nucleated cells counted after treatment with 10 μM of RP or ROL. H&E staining of sections was quantified using ImageJ software (average of 2 repeats, n=2 replicates each, \*\*\**p*<0.001). Scale bar (black) is 50 μm. D and E, Immunological analysis of proliferating cells within the epidermal equivalent after treatment with 10 μM ROL or RP. Cultures were immuno-stained with the proliferation marker Ki67 (green), the terminal differentiation marker filaggrin (red), and the DNA stain DAPI (blue). The number of Ki67 positive cells was quantified using ImageJ software (average of 2 repeats, n=2 replicates each, n=2, \*\*\**p*<0.001). Scale bar (white) is 50 μm.

## DISCUSSION

Retinoids are a broad family of molecules, several of which that include tRA, ROL, and RPalm have been utilized for topical treatment to improve the appearance of photoaged skin.^[30]^ More recently, RP has been found to clinically impact photoaged skin with minimal irritation.^[23-25]^ Since there have been no reports analysing a direct comparison between RP and the more established retinoids ROL and RPalm, we used a variety of *in vitro* and *ex vivo* skin models to measure their skin penetration, metabolic fate, impact on gene expression responsiveness and HAS activity as well as morphological changes in skin structure in 3D epidermal skin equivalents. In this study, metabolism of these retinoids showed a unique profile for RP compared to ROL and RPalm (Figure 4). RP topically applied to *ex vivo* skin biopsies, showed significant levels of the parent compound as well as the ROL metabolite in the viable epidermis and dermis, presumably due to rate and extent of retinyl ester hydrolysis.^[31]^ In contrast, ROL was primarily metabolized to RPalm and the inactive 14-HRR analogue. This confirms what has been previously reported that retinoid metabolites extracted from epidermis of human skin treated with 0.3% ROL treatment included ROL, 14-HRR, and retinyl esters.^[32]^ RPalm applied topically reached the viable epidermis and dermis to a lesser extent and remained there as the parent compound. It is not clear at present why RP and ROL have such different metabolite profiles. It’s possible that the rate of hydrolysis of the propionate ester group on RP provides for more parent compound to be maintained in the skin after 24 hours, while at the same time providing a rich source of ROL. Additionally, it’s possible that re-esterification of ROL to RPalm and/or metabolism to 14-HRR at longer time points beyond 24 hours.

**Figure 4.**
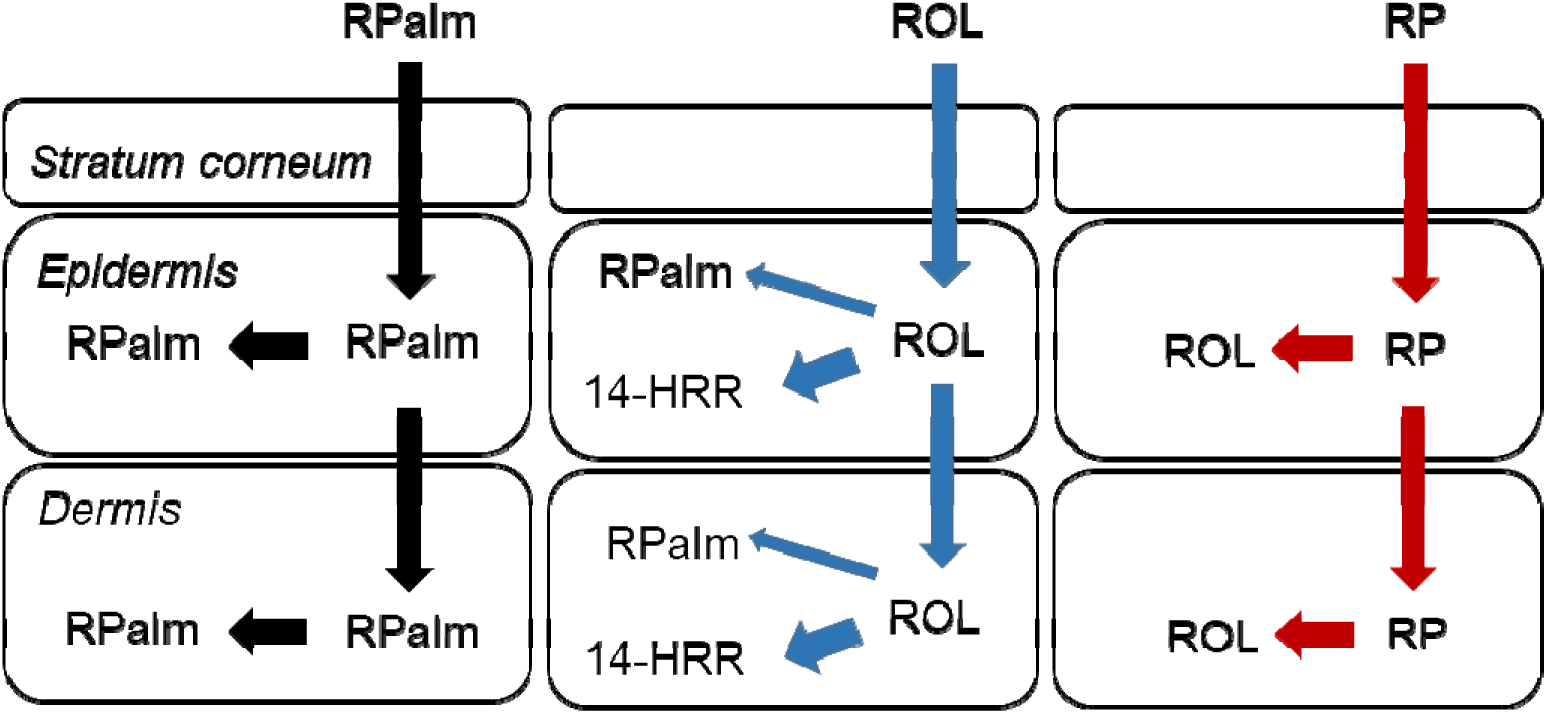
Schematic diagram of skin penetration and metabolism of RP, ROL, or RPalm. RP metabolic fate in viable skin is distinct from ROL and RPalm, showing preferred metabolism to ROL and no detectable esterification to RPalm or conversion to 14-HRR.

Further work is needed to better understand why RP has this unique advantage over ROL and RPalm. It is also interesting that we found that the majority of the retinoid applied did not penetrate and was removed after topical treatment of the skin biopsies with the retinoid formulations. Since we know that these retinoids, particularly RP, ROL, and tRA at these concentrations can deliver significant efficacy, it does present an opportunity to optimize the formulations such that lower levels of retinoids could be used while maintaining efficacy.

When keratinocytes were treated with 100 nM RP or ROL we measured a highly similar overall gene expression response pattern when comparing fold changes (r=0.98). Interestingly, PCR quantitation of a panel of 48 retinoid responsive genes identified several instances such as ELF3 and HBEGF where RP triggered higher expression levels than ROL in a dose dependent manner. Since its known that retinoids such as ROL and related esters can be metabolized by keratinocytes to tRA^[33]^, we utilized an RARα reporter cell line to measure the effects on an activated receptor mediated response. Both RP and ROL elicited an RARα activation response in a dose dependent manner. Again, RP showed an advantage over ROL for RARα activation at stoichiometrically equivalent concentrations across the tested doses. In contrast, RPalm did not show any retinoid activity in this model and in fact had an inhibitory effect at higher concentrations. It should be noted that HEK293 cells used to test for RARα activation are embryonic kidney cells and the results should not be extrapolated directly to how these retinoids may activate in keratinocytes. Thus, to further understand the differences between these retinoids, we also measured HA synthesis levels in proliferating keratinocytes since the enzymes encoding for hyaluronic acid synthase are retinoid regulated^[29]^ and found that HAS3 was upregulated by RP and ROL (Supplementary Table 2). RP showed a more consistent dose response on increasing HA synthesis over ROL and, again, was significantly higher across all concentrations tested compared to the equivalent concentration of ROL tested. In this model RPalm did increase HA levels compared to vehicle control but this response was significantly lower when compared to the lowest tested concentration of RP or ROL. These findings suggest that RP can elicit an overall retinoid response better than ROL and RPalm at the concentrations tested. This differential may be due to the unique metabolic fate of RP as measured in the *ex vivo* skin biopsies. Retinyl esters hydrolases metabolize retinyl esters into free ROL and these enzymes have been well studied in the dietary metabolism and systemic circulation of retinoids^[34]^. However, much less is known about REH enzymes in skin. While a putative REH has been reported in keratinocytes^[35]^, there has been limited research published since. It’s believed that RP is hydrolysed to ROL which in turn is oxidized to RAL and, ultimately, to tRA to elicit the measured retinoid responses. While ROL from either the parent compound or hydrolysed from RP or RPalm can be esterified by acyltransferases such as lecithin:retinol acyltransferase^[22]^, it is possible that the shorter alkyl chain length of RP in contrast to RPalm renders it having an altered recognition and binding affinity by REHs, acyltransferases, or other esterases. Alternatively, the kinetics of RP metabolism may be sufficiently different that we were not able to measure in the time frame of our experiments. Further work is needed to study the biochemistry and enzyme kinetics involved in the metabolite profiles between RP, ROL, and RPalm in these models.

Mechanistically, retinoids can restore epidermal thickening in photoaged skin by stimulating proliferation and modulating differentiation.^[17]^ Based on this, we utilized 3D epidermal skin equivalents as a novel skin model with a viable epidermis to evaluate what impact RP and ROL would have on the model’s morphology. Both RP and ROL at 10 μM caused a significant increase in epidermal thickness as well as increased staining for Ki67, a proliferation marker. Again, we found a significantly greater effect by RP over ROL at the concentration tested. Future work will utilize 3D epidermal skin equivalents to better understand the differences between these retinoids.

In conclusion, our collective data provides a body of evidence that RP can dose dependently elicit a stronger response than ROL and RPalm in retinoid sensitive *in vitro* and *ex vivo* models. We hypothesize that its unique metabolism profile, in part, provides this advantage over ROL and RPalm.

## Abbreviation list

RP: retinyl propionate
ROL: retinol
RPalm: retinyl palmitate
tRA: all-*trans*-retinoic acid
RARα: retinoic acid receptor-alpha
HA: hyaluronic acid
HAS: hyaluronic acid synthase

## DATA AVAILABILITY STATEMENT

The data reported in this paper have been deposited in the Gene Expression Omnibus (GEO) database, https://www.ncbi.nlm.nih.gov/geo (accession no. GSE155789). No other datasets were generated or analysed for this manuscript.

## AUTHOR CONTRIBUTIONS

J.E.O, D.L.B., R.L., J.D.S., K.G., V.M., S.A.P. designed the research. J.M.P., R.L.M.D., J.D.S., P.B.S., M.R., C.T., L.V., K.G., V.M. performed the research. J.E.O., D.L.B., R.L.M.D., R.L., C.T., M.R., K.G., V.M. analysed the data. J.E.O., D.L.B., J.D.S., K.G., V.M. wrote the paper. P.B.S., M.R., C.T., R.L.M.D., edited the paper. All authors have read and approved the final manuscript.

## ACKNOWLEDGEMENTS

We thank Charles River Laboratories Edinburgh Ltd for execution of the OECD 428 dermal penetration and metabolism study, Larry Robinson for retinoid formulation, and Holly Rovito for assistance with keratinocyte PCR experiments.

