## Supplemental File for "The vitamin A ester retinyl propionate has a unique metabolic profile and higher retinoid-related bioactivity over retinol and retinyl palmitate in human skin models"

**Supplemental material**

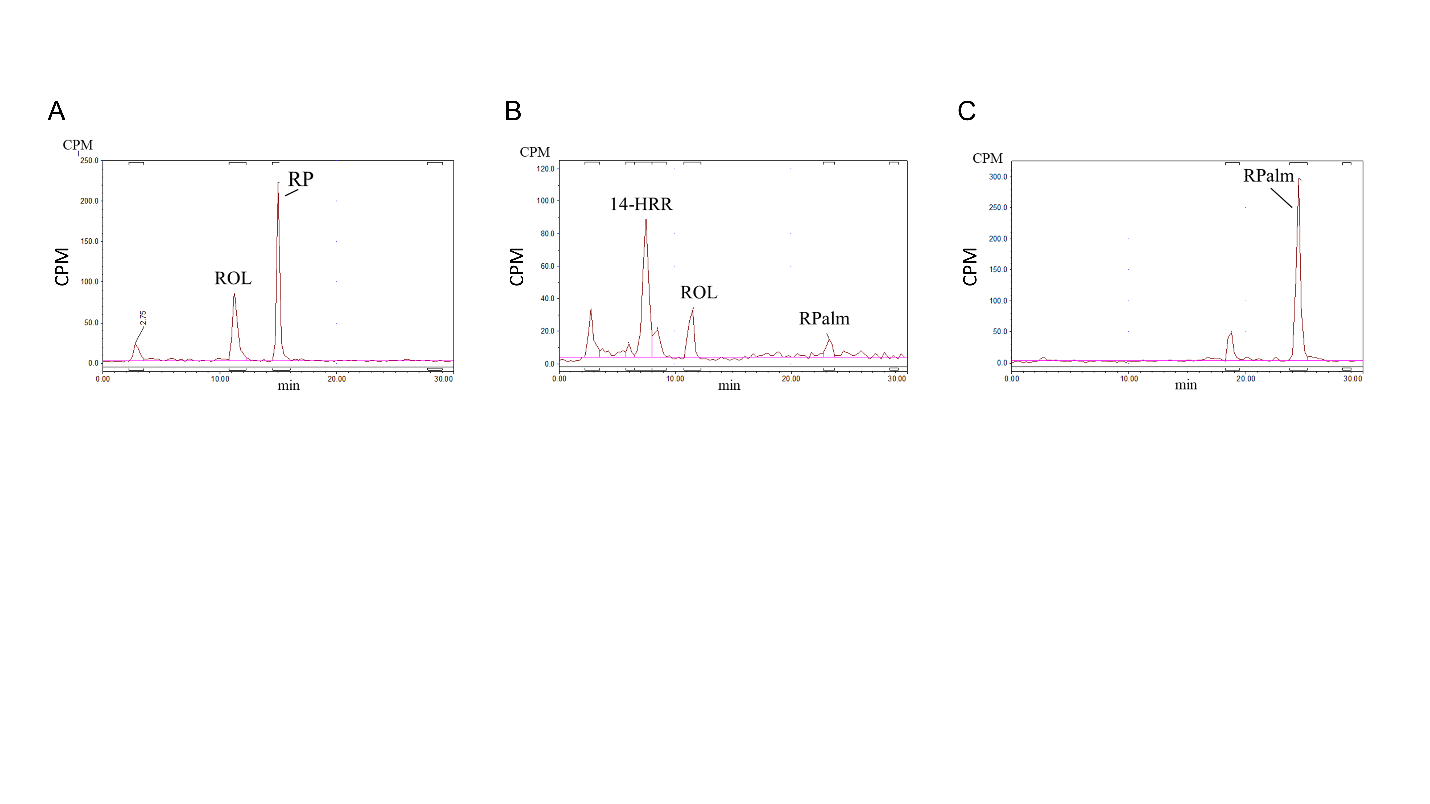

**Supplemental Figure 1.** Representative HPLC-RAD profiles of extracted ethanol-extracted epidermis, 24 hours following topical treatment with ^14^C-labeled: A, RP, B, ROL, C, RPalm. Parent compound and metabolite peak areas enable calculation of the relative proportion of significant ^14^C-labeled chemical species recovered from for each retinoid treatment.
